# Phylogenomic analysis reveals the evolutionary origins of five independent clades of forage grasses within the African genus *Urochloa*

**DOI:** 10.1101/2023.07.03.547487

**Authors:** Lizo E. Masters, Paulina Tomaszewska, Trude Schwarzacher, Alexandre R. Zuntini, Pat Heslop-Harrison, Maria S. Vorontsova

## Abstract

**Background and Aims:** The grass genus *Urochloa* (*Brachiaria*) includes forage crops that are important for beef and dairy industries in tropical and sub-tropical Africa, South America, and Oceania/Australia. Economically important species include *U. brizantha*, *U. decumbens*, *U. humidicola*, *U. mutica*, *U. arrecta*, *U. trichopus*, *U. mosambicensis*, and *M. maximus*, all native to the African continent. Perennial growth habits, large, fast growing palatable leaves, intra- and interspecific morphological variability, apomictic reproductive systems, and frequent polyploidy are widely shared within the genus. The combination of these traits likely favoured the selection for forage domestication and weediness, but trait emergence across *Urochloa* cannot be modelled, as a robust phylogenetic assessment of the genus has not been conducted.

**Methods:** Using a target enrichment sequencing approach (Angiosperms353 baits), we inferred a species level phylogeny for *Urochloa sensu lato*, encompassing 57 species (∼50% of the genus) and outgroups. We determined the phylogenetic placement of agriculturally important species and identify their closest wild relatives. Further, we mapped key traits associated with forage crop potential to the species tree, exploring trait distribution across the genus.

**Key Results:** Agricultural species belong to five independent clades, including *U. brizantha* and *U. decumbens* lying in a previously defined species complex. Crop wild relatives were identified for these clades supporting previous sub-generic groupings in *Urochloa* based on morphology. Using ancestral trait estimation models, we find that five morphological traits that correlate with forage potential (perennial growth habits, culm height, leaf size, a winged rachis, and large seeds) independently evolved in forage clades.

**Conclusions:** *Urochloa s.l.* is a highly diverse genus that contains numerous species with agricultural potential, including crop wild relatives that are currently underexploited. The African continent is the centre of origin for these clades and their conservation across their native distributions is essential. Genomic and phenotypic diversity in forage clade species and their wild relatives needs to be better assessed to improve sustainability in *Urochloa* cultivar production.

## Introduction

African grasses have been recognised for their forage potential since the 18^th^ century, and as a result have been transplanted around the globe to upscale beef and dairy production for small-scale and commercial farms (Hartley and Williams, 1956; Parson, 1976; Cook and Dias, 2006; Visser et al., 2016). Today, arguably the most important of these African grasses belong to the genus *Urochloa* P. Beauv., a large and diverse genus including taxa previously placed in *Brachiaria* (Trin.) Briseb., *Chaetium* Nees, *Eriochloa* Kunth, *Scutachne* Hitchc. & Chase, and *Megathyrsus* (Pilg.) B.K. Simon & S.W.L. Jacobs (Salariato et al., 2010; Salariato et al., 2012; Jank et al., 2014; Kellogg, 2015; Namazzi et al., 2020, Ferreira et al., 2021). *Urochloa* forages are strongly preferred in sub-tropical and tropical regions as they are highly palatable and nutrient dense, are tolerant of low-quality soils, and outcompete alternative forage grasses in terms of biomass productivity e.g., *Pennisetum purpureum* Schumach. and *Cenchrus ciliaris* Fig. & De Not. (Maass et al., 2015; Baptistella et al., 2020, Njarui et al., 2021; Rathore et al. 2022). Since the 1950s these grasses have been adopted in forage systems in Southeast Asia, Australia, and especially Central and South America, with an estimated 99 million hectares of land devoted to *Urochloa* production in Brazil alone (ANUALPEC, 2008; Jank et al., 2014).

Livestock rearing for meat and dairy are key areas of economic importance in numerous developing nations, contributing greatly to the livelihoods of rural communities (e.g. Colombia) and commercial farmers (e.g. in Brazil) (Jank et al., 2014; Enciso et al., 2021). Recent breeding projects lead by the Centro Internacionale de Agricultura Tropical (CIAT; now Alliance Bioversity & CIAT), Colombia, and Empresa Brasileira de Pesquisa Agropecuária (EMBRAPA), Brazil, have produced highly productive and globally competitive *Urochloa* cultivars (Miles and Valle, 1996; Valle and Savidan, 1996; do Valle et al., 2013; Worthington and Miles, 2015; Espitia et al., 2020). High demand has resulted in the reintroduction of *Urochloa* cultivars into African countries including Kenya, Uganda, and Madagascar, largely for improving milk production in small-holder farms (Maass et al., 2015; Mutimura et al. 2018).

Cattle rearing contributes greatly to global methane emissions, and the conversion of natural habitats to agricultural forage lands has contributed to global biodiversity loss (Herero et al., 2011; Peters et al., 2012; Godfray et al., 2018). African *Urochloa* species may be invasive outside of their natural ranges, recognized in Australia and the Americas (Foxcroft et al., 2010; Seabloom et al., 2013; Visser et al., 2016; Overhalt and Franck, 2017). African *Urochloas* aggressively exclude indigenous vegetation in disturbed environments through a combination of rapid growth and spread, and the production of allelopathic chemicals which hinder the development indigenous flora (Barbosa et al., 2008; Kato-Noguchi et al., 2014; Damasceno et al., 2018). Further, *Urochloa* forages possess undesirable traits, chiefly an inability to survive frost and are typically grown in monocultures in large-scale systems (do Valle et al., 2013; Krahl et al., 2019). Although recent decades have seen tremendous advances in *Urochloa* breeding, challenges in understanding the taxonomy and evolutionary history in *Urochloa* species remain and limit breeders to a relatively small pool of taxa and accessions to choose from, slowing down breeding (do Valle et al., 2013; Sweitzer et al., 2021). As one of the most economically significant forage genera across the tropics, improving the sustainability of *Urochloa* cultivars is paramount for achieving development goals.

*Urochloa* cultivar breeding is difficult due to asexual reproduction via apomixis and diverse ploidy levels present in most genotypes grown as forages (Miles and Valle, 1996; Miles 2007; Worthington and Miles, 2015; Worthington et al., 2016; Hanley et al., 2021; Tomaszewska et al., 2021). To overcome the challenges posed by apomixis and polyploidy, forage breeders artificially induce polyploidy in sexually reproducing diploids and cross them with closely related apomictic polyploid species (Ishigaki *et al*. 2009). For example, tetraploids of the apomictic species *U. brizantha* and *U. decumbens* are crossed with artificial tetraploids of the sexual diploid *U. ruziziensis,* where apomictic acted as pollen donors (Miles and do Valle, 1996). Crosses of these three species have produced the most commercially successful *Urochloa* cultivars (Pizarro *et al*. 2013). Exploiting interspecific hybridization is central to modern *Urochloa* forage breeding, and successful cultivar production requires in-depth knowledge of the taxonomy and ploidy of the species used for crossing. In *U. brizantha*, *U. decumbens*, and *U. ruziziensis*, molecular data have revealed their shared, complex evolutionary history. While all three species are clearly closely related, *U. brizantha* and *U. ruziziensis* belong to divergent lineages with parts of *U. decumbens* divided between the two by ploidy levels (tetraploids grouping with *U. brizantha* and diploids with *U. ruziziensis*) (Triviño et al., 2017; Higgins et al., 2022; Tomaszewska et al., 2023).

Taxonomic uncertainty, mosaics of sexually and asexually reproducing close relatives, and diverse intraspecies ploidy levels are typical of *Urochloa* forage species, and forage grasses more generally (Sandhu et al., 2019; Ortiz et al., 2020). This is the case for the largely apomictic, hexaploid species *U. humidicola*, the third most agriculturally significant *Urochloa* crop species after *U. brizantha* and *U. decumbens* (Vigna et al., 2016; Worthington et al., 2019). *Urochloa humidicola* populations include intermediates morphologically more similar to *U. dictyoneura*, leading taxonomists to argue that the two species should be lumped into one (Sosef, 2016). Two additional agricultural species complexes (containing *U. mutica* with *U. arrecta*, and *U. trichopus* with *U. mosambicensis* respectively) can be characterised by overlapping morphologies, and apomictic reproduction (Toutain, 1986; Pereira Filho et al., 2013). A further complication is that the modern taxonomic concept of *Urochloa*, a monophyletic clade comprising previously separate genera, now includes *Megathyrsus maximus* (synonyms *Panicum maximum* Jacq. and *U. maxima* (Jacq.) R.D. Webster), a globally significant tetraploid forage and weed species indigenous to the African continent (Salariato et al., 2012; Rhodes et al., 2021).

Although belonging to different species complexes, these important forages share numerous agriculturally relevant traits (Keller-grein et al., 1996). *Urochloa* forage species display perennial growth habits, a trait directly associated with carbon sequestration (Lal, 2004; Wilson et al., 2018; Ledo et al., 2020). Compared to their wild relatives, *Urochloa* forage grasses are generally characterised by their tall culms, large and broad leaves, and produce relatively large seeds (Fisher and Kerridge, 1996; Clayton et al., 2016). Grasses which naturally produce high biomass yields through leaves and culms are desirable in tropical forage systems, while the production of large and easily harvested seeds is an advantageous trait for seed multiplication (Juntasin *et al*. 2022). *Urochloa* cultivars are predominantly sold and distributed through seed, and high yields in vegetative biomass and easy multiplication through seed are a dual aim for forage breeders (Hopkinson et al., 1996; Santos Filho, 1996, Ghimire et al., 2015). The inflorescences of forage species typically consist of simple, unbranched racemes (Reinheimer and Vegetti, 2008; Salariato et al., 2010), and with laterally elongated or ‘winged’ rachises (Clayton and Renvoize, 1982; Renvoize et al., 1996). These specific traits are not only agriculturally significant, but they are also directly measurable in wild *Urochloa* species through herbarium collections, or they are recorded in taxonomic treatments and floras (Clayton and Renvoize, 1982). Thus, modelling the evolution of these forage traits across *Urochloa s.l.* is possible and can provide insight into the emergence of wild species with forage potential. To achieve this, a comprehensive and robust phylogeny for *Urochloa* must be defined.

Phylogenetic studies of *Urochloa* and all subsumed genera (hereafter *Urochloa sensu lato*) have been limited to only a select few species, focusing largely on relationships between known economically important species i.e., *U. brizantha*, *U. decumbens*, and *U. humidicola* (González and Morton, 2005; Pessoa-Filho et al., 2017), or have sampled the genus broadly but inferred phylogenetic placements using a handful of chloroplast genes and/or nuclear barcoding markers (Salariato et al., 2010; Washburn et al., 2015; Hackel et al., 2018). Understanding the evolution of important forage species in *Urochloa s.l.* requires a phylogeny inferred across a broad representation of the genus using a large set of independently evolving gene regions. Modelling the evolutionary history of *Urochloa* with such a dataset will allow us to accurately infer speciation events in the face of population dynamics such an incomplete lineage sorting (Maddison 1997; Edwards 2009), account for gene duplications because of polyploidization (Wendel, 2015), and provide more accurate estimates for the emergence of species with forage crop potential. Wild *Urochloa* species that share morphologies with forage species have been identified (Renvoize et al., 1996), but to our knowledge these relationships have not been phylogenetically tested.

We aim to infer a genus wide phylogeny (species tree) for *Urochloa s.l.* using the Angiosperm 353 target enrichment of nuclear genes (Johnson et al., 2019) and infer the placement of the following agriculturally important species: *U. brizantha, U. decumbens*, *U. ruzizie*nsis, *U. humidicola*, *U. mutica, U. arrecta*, *U. trichopus*, *U. mosambicensis*, and *M. maximus*. As crop wild relatives (CWR) (Harlan and De Wet, 1971) for *Urochloa* forages, we aimed to determine which *Urochloa* species are most closely related to those used as forage species, based on most recent common ancestors. Finally, we infer ancestral trait estimates along a phylogeny for the following agriculturally important traits: perennial vs annual life cycle, culm height, leaf size, rachis wing morphology, and seed size. This will allow us to estimate the emergence of species with forage potential across *Urochloa s.l*.

## Materials and Methods

### Taxon sampling

A total of 64 samples representing 61 species were chosen for phylogenomic analysis using the Angiosperm353 target capture probe set (Johnson et al., 2019). The complete dataset contained 57 species within *Urochloa s.l.* including *Brachiaria* species within the Boivinellinae clade (Hackel et al., 2018), 5 additional species within the Melinidinae subtribe, and *Anthephora hermaphrodita* (L.) Kuntze and *Poecilostachys oplismenoides* (Hack.) Clayton as outgroup taxa within the Paniceae tribe (see Kellogg 2015 and Soreng et al., 2017 for classification within Poaceae). Taxa were chosen to meet two objectives: Sample species broadly across *Urochloa s.l.* and to target putative CWRs. The Kew Herbarium index, which orders species within genera based on morphological similarities, Renvoize and Maass (1993), and Renvoize et al. (1996) were used as guides for putative CWR sampling. A summary of all samples used with metadata for downstream analysis and data sources can be found in supplementary table 1. Leaf material from herbarium and silica dried specimens was then used for DNA extraction and target enrichment library preparation.

### Extraction and Target-enrichment Sequencing

#### DNA extraction, library preparation and sequencing

DNA extraction was performed using a modified CTAB protocol (Doyle and Doyle, 1987). Approximately 20mg of dried leaf material per sample was crushed into a fine powder and incubated in 750µl of Isolation Buffer (747µl CTAB with 3µl of 2-Mercaptoethanol) at 65°C for 30 minutes. 750µl of Chloroform:Isoamyl alcohol (24:1) was added to all samples which were then incubated at 65°C for 30 minutes in an orbital shaker set to 250 rev/mins. Following incubation, samples were centrifuged for 15 minutes at 13000rpm. 600µl of the DNA-CTAB complex was removed after centrifugation and placed in microcentrifuge tubes with 500µl of Isopropanol. Samples were homogenised and placed in -20°C freezer for 1 week. Following this period, samples were centrifuged for 15 minutes at 13000rpm. Supernatants were discarded and the pellet for all samples were washed with 750µl of 70% ethanol for 15 minutes. The supernatant was again discarded, and samples were left to air dry overnight before being resuspended in TE buffer. Gel electrophoresis was conducted to estimate extraction quality and DNA fragment size. Library preparation was conducted following the Illumina NEBNext Ultra II kit protocol, performing half volume reactions. Library concentrations were normalised to reduce variation across all samples, and samples were pooled into shared tubes for multiplex sequencing based on average fragment lengths. Multiplexed samples were then sent to Macrogen Inc., South Korea for target-enrichment sequencing.

### Phylogenomic Inference

#### Read processing and loci assembly

Paired end and unpaired raw reads for newly sequenced samples and samples obtained from various DNA sequence read archives were trimmed of adapters and filtered for low quality using Trimmomatic 0.39 (Bolger et al., 2014). Quality parameters were set using a Phred33 score. A minimum read length was set to 36 base pairs. Trimming was performed by assessing the leading and trailing read ends, and a sliding window of 4 base pairs was used across reads. Base pairs falling below the quality threshold were removed.

Trimmed reads were used for loci assembly using HybPiper 1.3.1 (Johnson et al., 2016). An amino acid target fasta file was used to capture on-target reads for Angiosperm 353 loci. This was to ensure synonymous mutations did not bias read mapping to the target files across divergent taxa. Loci were then assembled and retrieved for each sample using the ‘reads_first.py’ and ‘retrieve_sequences.py’ scripts from HybPiper. Only the exons of assembled genes were used for downstream phylogenomic analysis. This was chosen to standardise alignments in the dataset which contained both target-enrichment and transcriptomic sequences.

#### Paralogue removal and consensus sequence inference

In target-enrichment based phylogenomic inference, paralogous genes are commonly removed from the analysis as they introduce homoplasy and confound estimations of species divergence histories (Nicholls et al., 2015; Andermann et al., 2020; Larridon et al., 2020, Crowl et al., 2022). Further, the presence of paralogues and duplicated genes can confound gene assembly, as reads from different paralogues can be adjointed into contigs (and subsequently, genes), leading to chimeric sequence assembly (Kates et al., 2018; Nauheimer et al., 2021). The tradeoff to removing entire genes with suspected paralogy is that genes with high allelic diversity (i.e. in the case of whole genome duplication and reticulation events) can be purged, resulting in the loss of informative, non-paralogous loci (Morales-Briones et al., 2022). Whole genome duplications and interspecific reticulation events are ubiquitous in angiosperms, and specifically prevalent in Poaceae and *Urochloa* (Van der Peer et al., 2009; Vigna et al., 2016; McKain et al., 2016; Landis et al., 2017).

To strike a balance between paralogue removal and diverse orthologue retention, we assessed heterozygous loci with high allele diversity using SNP assessment scripts in HybPhaser (Nauheimer et al., 2021). We assessed SNP diversity for all 353 loci in all samples and removed loci with a SNP diversity that was 1.5x greater than the third interquartile range for SNP diversity in the entire gene dataset. This heuristic method allowed us to assess the relative heterozygosity across *Urochloa* loci and flag outlier genes as potential paralogous, while retaining duplicated orthologues resulting from polyploidy events. This resulted in the removal of 26 putatively paralogous genes leaving 327 loci for downstream analysis.

To account for gene variants in putative non-paralogous genes, we inferred ambiguously coded consensus sequences for each gene using a Phasing Alleles from Target Enrichment data (PATE) pipeline (Tiley et al., 2021). Estimated ploidy levels for all samples is required for the pipeline and were inferred from literature (Morrone et al., 2006; Tomaszewska et al., 2021), the Chromosome Count Database (http://ccdb.tau.ac.il/), or using the estimatePloidy.pl script in PATE. Ploidy levels are estimated by mapping reads to a reference sequence, in this case loci from the known diploid *U. fusca* (Morrone et al., 2006). Samples are initially genotyped as diploids using GATK3.8.1 (McKenna et al., 2010) and biallelic reads were then mapped to *U. fusca* loci. The ratio between reads matching the reference sequence and reads carrying the alternate were used to estimate ploidy levels (Tiley et al., 2018). For ploidy estimation, a maximum ploidy level of 6 was chosen to constrain the analysis.

Following ploidy estimation, samples were phased for gene variants with the maximum number of possible haplotypes determined by the estimated ploidy levels. Reads per gene per sample were assigned to sequences from HybPiper using BWA 0.7.17 (Li and Durbin, 2009) and PCR duplicates were flagged using Picard v2.27.4 (http://broadinstitute.github.io/picard). HaplotypeCaller in GATK3.8.1 was used to assign reads to gene variants with parameter settings left as default in the PATE.pl script (Tiley et al., 2021). Phased gene variants were then assembled using H-PoPG v0.2.0 (Xie et al., 2016). Phased variants were then collapsed into consensus sequences where polymorphic sites were coded with ‘N’ to ensure that chimeric assembly of homoeologous sequences were not present in phylogenetic analysis.

#### Species tree inference

Ambiguously coded sequences were aligned using MAFFT v7.475 (Katoh and Standley, 2013) with parameters set to L-INS-I for the highest stringency. Columns in alignments with more than 30% missing data were removed using Phyutility v2.2.6. (Smith and Dunn, 2008) Individual maximum likelihood genes trees were inferred using IQTREE v2.1.2 (Minh et al., 2022) with ModelFinder Plus (Kalyaanamoorthy et al., 2017) used to determine the best fit model per gene based on BIC scores. Ultrafast bootstrapping was implemented to assess node support using 1000 replicates (Hoang et al., 2018). Gene trees were then concatenated into a single file and nodes with support values of 10 or less were collapsed using Newick Utilities v1.6 (Junier and Zdobnov, 2010). Outlier taxa with excessively long branch lengths were then removed from all gene trees using TreeShrink (Mai and Mirarab, 2018). A species tree was then inferred using ASTRAL-III v5.7.7 (Zhang et al., 2018). Separately, gene alignments were concatenated into a supermatrix which was used to infer a maximum likelihood (ML) phylogeny using IQTREE v2.1.2. 1000 ultrafast bootstrap replicates were used to assess node support and a GTR+G+I nucleotide substitution model was chosen due to computational constraints.

### Character evolution

#### Trait data

For continuous traits we used maximum leaf area, maximum culm height, and maximum seed size (inferred from maximum fertile lemma length). For discrete characters we used growth habit (annual vs perennial growth habit) and rachis wing morphology (wingless, narrowly winged, or broadly winged). Trait data were taken from the GrassBase (Clayton et al., 2016). Data were filtered for relevant species and traits using the R packages tidyr (Wickham et al., 2023a) and dplyr (Wickham et al., 2023b). Updating and reconciling species names between our samples and the GrassBase database was done using the World Checklist of Vascular Plants (WCVP) (Govaerts et al., 2021), accessed through Plants of the World Online (https://powo.science.kew.org/), and Vorontsova (2022).

#### Ancestral estimation methods

Estimations for ancestral traits were inferred using the supermatrix ML tree. Taxa with no trait data and duplicate taxa were removed from the tree using the ‘drop.tip’ function in the R package ape (Paradis et al., 2004). For continuous traits (leaf area, culm height, and fertile lemma length), values were log transformed and maximum likelihood of ancestral states were estimated under a Brownian motion model using the fastANC function from the R package phytools (Revell, 2012). To account for uncertainty in trait estimations at nodes, the variance and confidence intervals for every node were calculated too (Losos, 1999). We then tested for phylogenetic signal, the tendency for closely related taxa to share trait values more frequently than by chance (Revell et al., 2008), in continuous traits using Blomberg’s K (Blomberg et al., 2003) and Pagel’s **λ** (Pagel 1999).

To determine the best fit model for growth habit we compared ‘Equal Rates’, ‘All Rates Different’, ‘Perennial to Annual’ (but not reversible), and ‘Annual to Perennial’ (but not reversible) models following Revell and Harmon (2022). The model with the best Akaike Information Criterion (AIC) score was chosen for analysis. The same approach was used for rachis wing morphology. The models compared were ‘Equal Rates’, ‘All Rates Different’, and ‘Symmetrical Rates’, following Revell and Harmon (2022). Ancestral state estimations were then inferred using a marginal likelihood ancestral state reconstruction method using the R package corHMM (Beaulieu et al., 2013). The posterior probabilities for ancestral state were then mapped as pie charts to internal nodes of the tree using the R package ggtree (Yu et al., 2017).

## Results

### Phylogeny and origins of forages

The ASTRAL-III and supermatrix ML trees recovered similar tree structure for *Urochloa* forage species. Congruence between the two topologies was used to define five clades giving rise to forages (figure 1, table 1). In both trees, *U. humidicola* and *U. dictyoneura* form a clade with the wild species *U. brevispicata*, *U. stigmatisata*, *U. reticulata,* and *U. dura*, hereafter referred to as the *‘Humidicola* clade’. The clade is well supported with 100% bootstrap values in both trees, and high gene tree congruence in the ASTRAL-III analysis. The three most commercially important *Urochloa* forages, *U. brizantha, U. decumbens*, and *U. ruziziensis* formed a clade with wild species *U. eminii* and *U. oligobrachiata* with 100% node support in both trees, forming the *‘Brizantha* clade’. *Urochloa arrecta* and *U. mutica* form a highly supported clade (*Mutica* clade) in both trees but the ASTRAL-III analysis placed them sister to *Scutachne dura*, a monotypic species from the Caribbean (Webster and Reyna, 1988), with low node support (71%) and gene tree congruence. The placement of the *Mutica* clade is also incongruent between the two tree analyses: ASTRAL-III places the clade sister to the *Brizantha* clade and *Humidicola* clade, where the ML tree shows the *Mutica* clade and *Brizantha* clade are closely related and the *Humidicola* clade belongs to a sister lineage.

**Figure 1:**
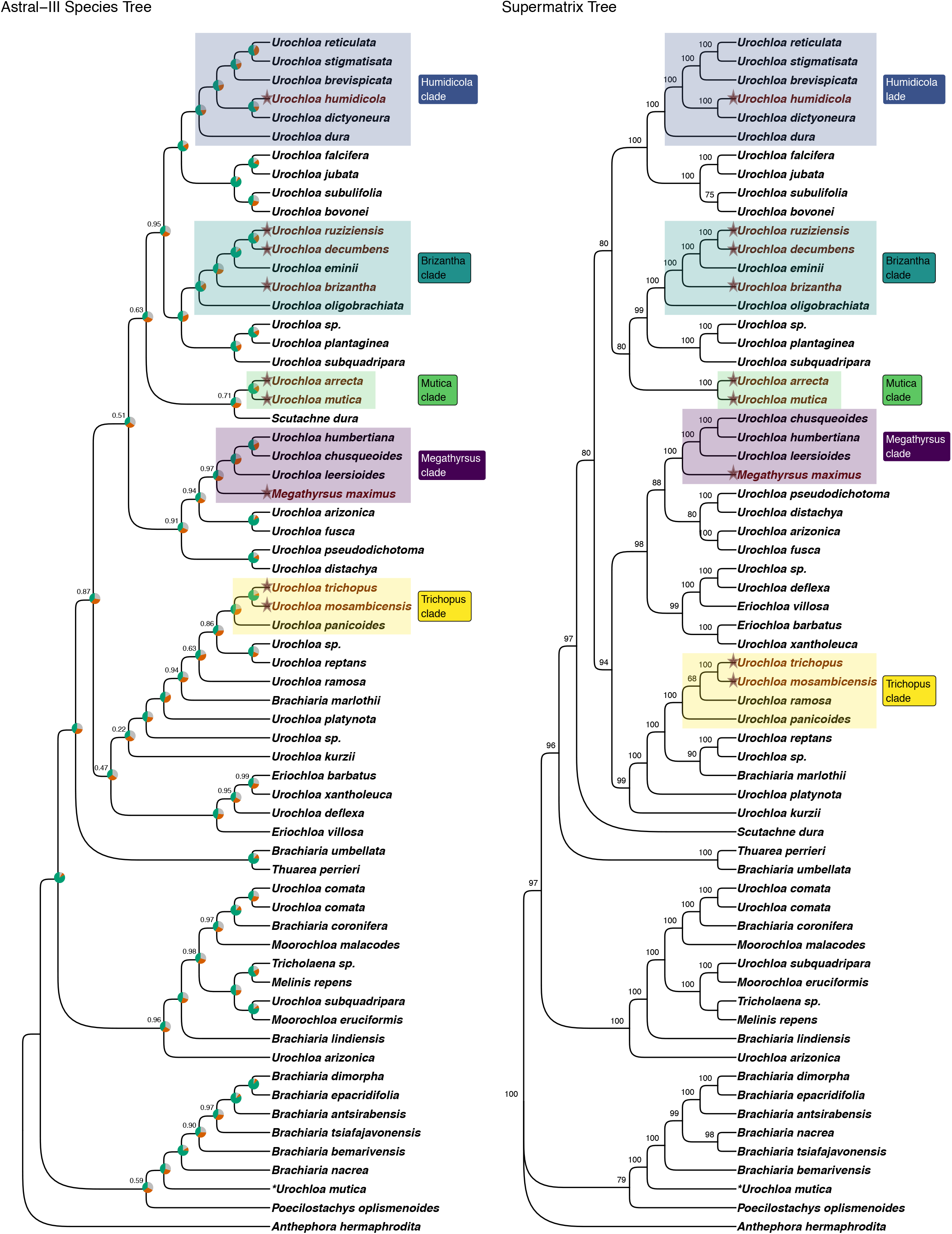
Phylogenetic reconstruction of the evolutionary history of *Urochloa sensu lato* using 327 nuclear markers. *Urochloa* forage species are marked with a star and labelled in red. Forage clades with CWRs are highlighted. (Left) Species tree of *Urochloa* inferred using the summary method in ASTRAL III. (Right) Phylogeny of *Urochloa* inferred from a concatenated DNA supermatrix using a maximum-likelihood method in IQTREEv2.1.2

**Figure 2:**
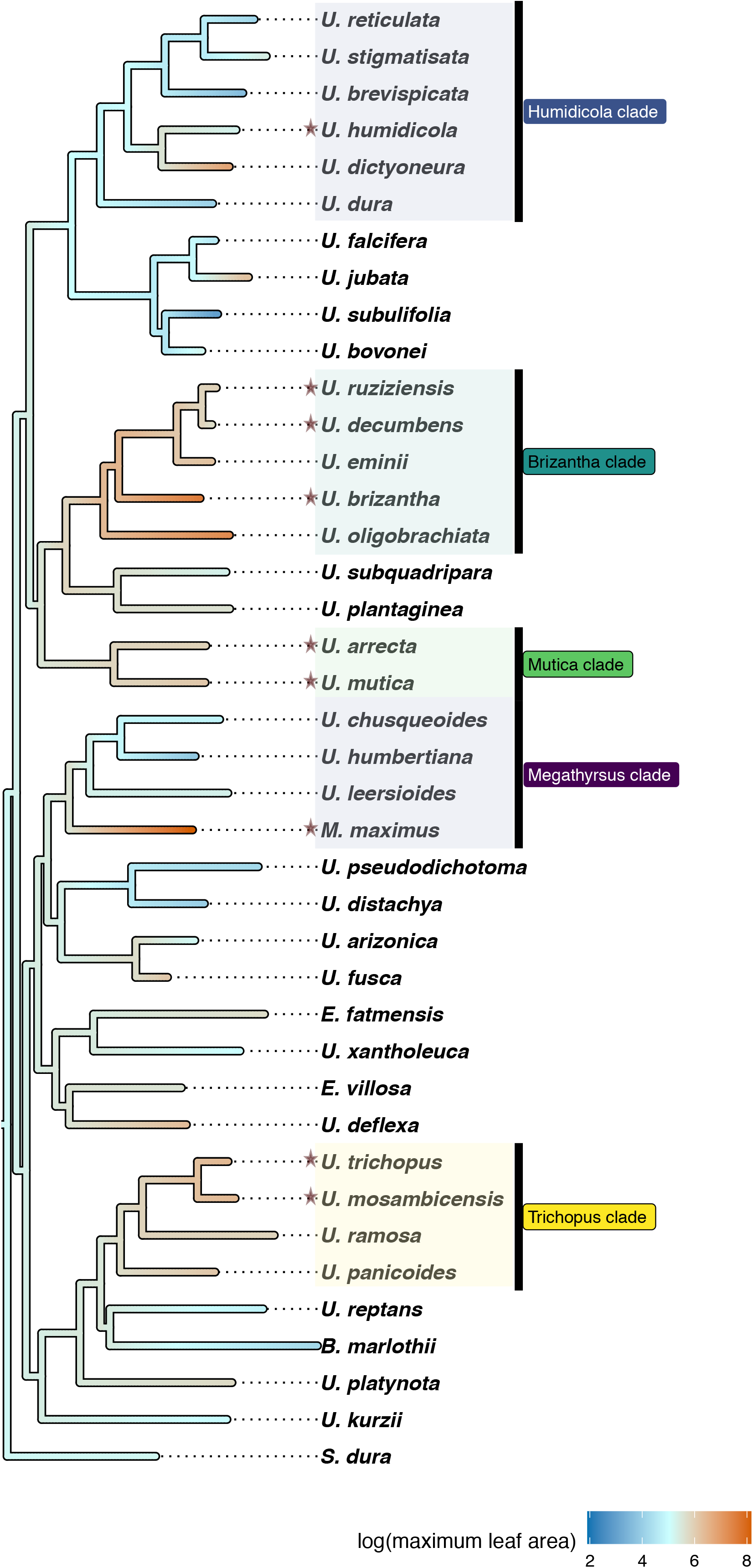
Evolution of leaf area (cm^2^) in *Urochloa s.l.* tree excluding Melinidinae, Boivinellinae and outgroup taxa. Forage species are marked with a star. Ancestral state estimations are inferred along branch lengths.

**Table 1:**
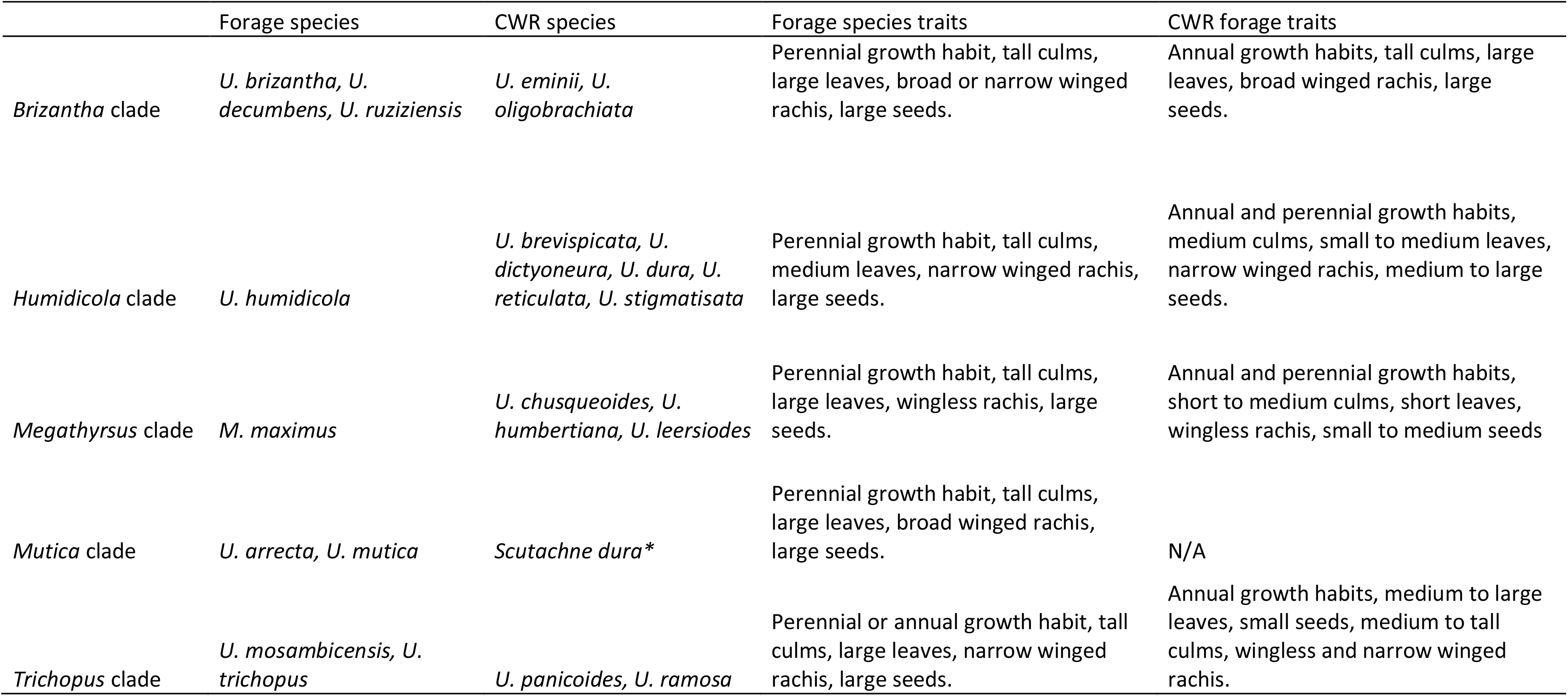
*Urochloa* forage crop clades and their crop wild relatives (CWR) defined here. Comparisons in forage traits between forage species and CWR are summarised. \**Scutachne dura* was only supported as a CWR to the *Mutica* clade in the ASTRAL III species tree not used for ancestral state estimation analyses.

Both trees confirmed the placement of *M. maximus* within *Urochloa* and placed the taxa within a clade containing close relative *U. humbertiana*, *U. leersioides*, and *U. chusqueoides* (*Megathyrsus* clade). Both trees have 100% bootstrap support for this clade, but ASTRAL-III analysis shows high gene tree incongruence at this node. *U. trichopus* and *U. mosambicensis* formed a highly supported clade containing the wild relative *U. panicoides* in both analyses (the *Trichopus* clade). However, the ML tree placed *U. ramosa* within this clade as well, and was more closely related to the forage species than *U. panicoides*. Apart from three forage species within the *Brizantha* clade, all forage species belonged to independent lineages within *Urochloa s.l.,* despite their shared geographical origins on the African continent.

In addition to the genera *Scutachne* and *Megathyrsus,* both trees support placing *Eriochloa* within *Urochloa,* and placing *Moorochloa* as a separate though polyphyletic genus. *B. umbellata* is closely related to the genus *Thuarea* (Hackel et al. 2018) and sits outside the *Urochloa* clade in our analysis. In both trees, our analysis supports the Boivinellinae clade, containing ‘*Brachiaria*’ that are endemic to Madagascar, as sister to the Melinidinae subtribe Hackel et al., 2018). Both trees show that *U. arizonica* and *U. subquadripara* are paraphyletic with a specimen from each species falling within the clade *Urochloa,* and a specimen placed outside the clade.

### Ancestral trait estimations

Ancestral states for the chosen continuous traits showed moderate size for leaf area, culm height, and seeds across not just *Urochloa*, but the Melinidinae subtribe and Malagasy *’Brachiaria’* species (leaf area figure 1; supplementary figures 1 - 3). For the *Brizantha* and *Mutica* clades, ancestral state estimations for log leaf area and log culm height show an increase in size of these traits at the node of their respective most recent ancestors. The *Humidicola*, *Megathyrsus*, and *Trichopus* clades show greater variability in these two traits, as agriculturally significant species differ from their closest relatives and their respective estimated ancestors. This is most observably clear for *M. maximus*, which evolved much larger leaves and culms than its closest living relatives and their most recent shared ancestor.

Estimates for log fertile lemma length show *M. maximus*, *U. trichopus*, *U. mosambicensis*, and *U. arrecta* evolved large seeds from small-seeded ancestors independently. Ancestral state for the *Humidicola* clade indicates that the common ancestor likely had moderately large seeds, those seed size increased along the *U. humidicola*/*U. dictyoneura* lineage and decreased in wild relatives. Long fertile lemma length estimates for the *Brizantha* clade indicate that this clade evolved from a common ancestor with large seeds. Phylogenetic signal for all three continuous traits was high and statistically significant, indicating that the similarity in trait values across all taxa is the result of a shared phylogenetic history (table 2.

**Table 2:** Phylogenetic signal values (Blomberg’s K and Pagel’s **λ**) with p-values for natural log values of continuous traits (leaf area, lemme length, and culm height).

| | Blomberg's K | p-value | Pagel's $\lambda$ | p-value |
| --- | --- | --- | --- | --- |
| log Leaf Area | 0.775067 | 0.0016 | 0.800593 | 9.51344e<sup>-5</sup> |
| log Lemma Length | 0.68713 | 0.0052 | 0.999934 | 0.000952218 |
| log Culm Height | 0.636892 | 0.0254 | 0.891587 | 0.00495678 |

AIC scores determined that an ‘Equal Traits’ model was the best fit for ancestral trait estimation for both discrete traits i.e. growth habit and rachis wing morphology (table 3). Posterior probability scores show that the ancestor of *Urochloa* grasses was likely an annual and that perennial growth habits likely emerged multiple times at ancestral nodes within the genus (figure 3). Estimations of ancestral rachis wing morphology strongly show that *Urochloa s.l.* evolved from an ancestor with a wingless rachis (figure 4). The emergence of a rachis wing (narrow or broad) occurred in parallel across multiple nodes in the phylogeny and is generally associated with forage clades (a notable exception being the *Megathyrsus* clade where all members are wingless).

**Figure 3:**
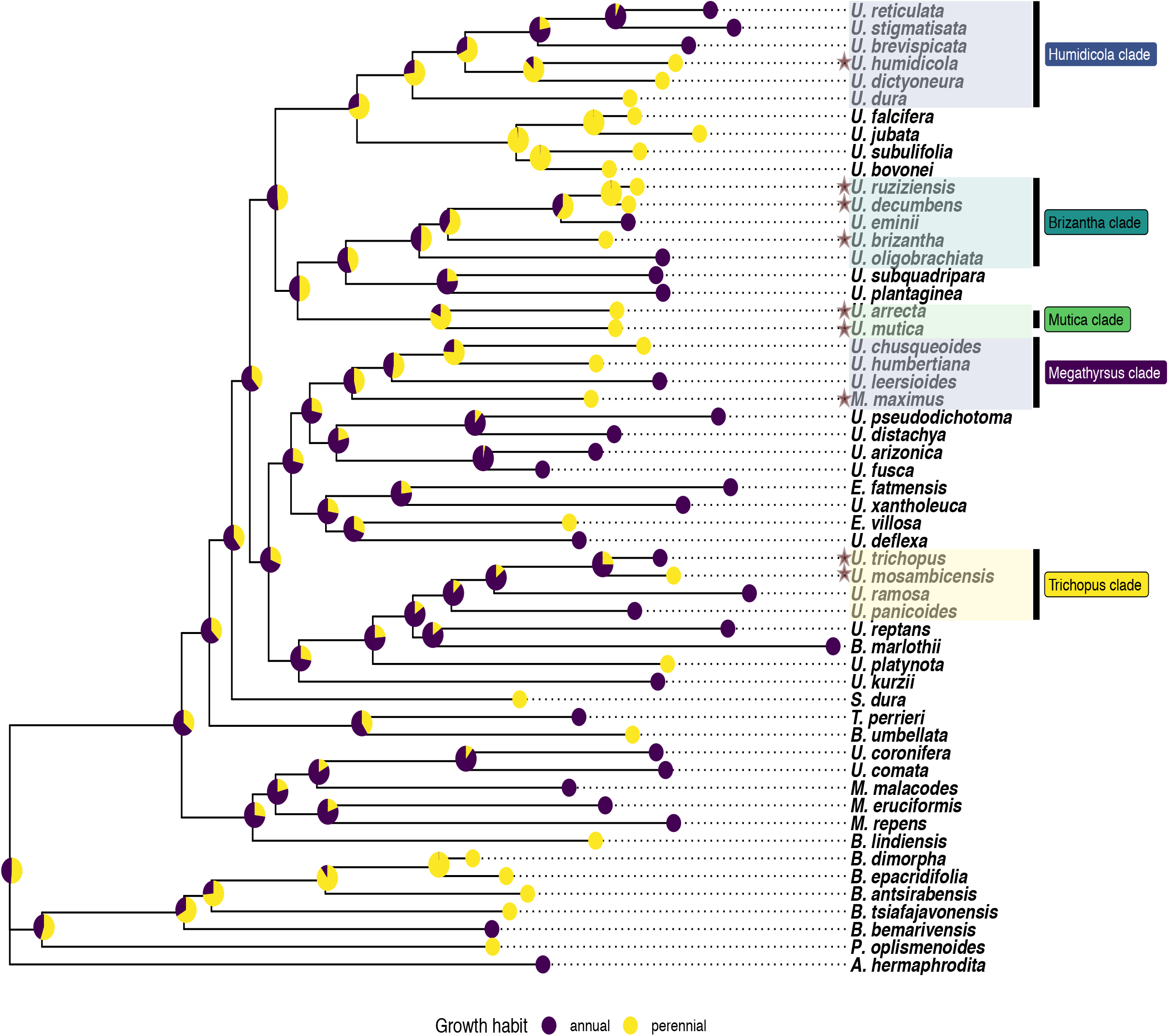
Evolution of growth form (annual vs perennial). Maximum likelihood tree estimated using IQTREE2 v2.1.2 inferred from 327 nuclear genes. Ancestral state estimations for annual versus perennial habits were performed using corHMM in R and posterior probabilities for state estimations mapped to ancestral nodes.

**Figure 4:**
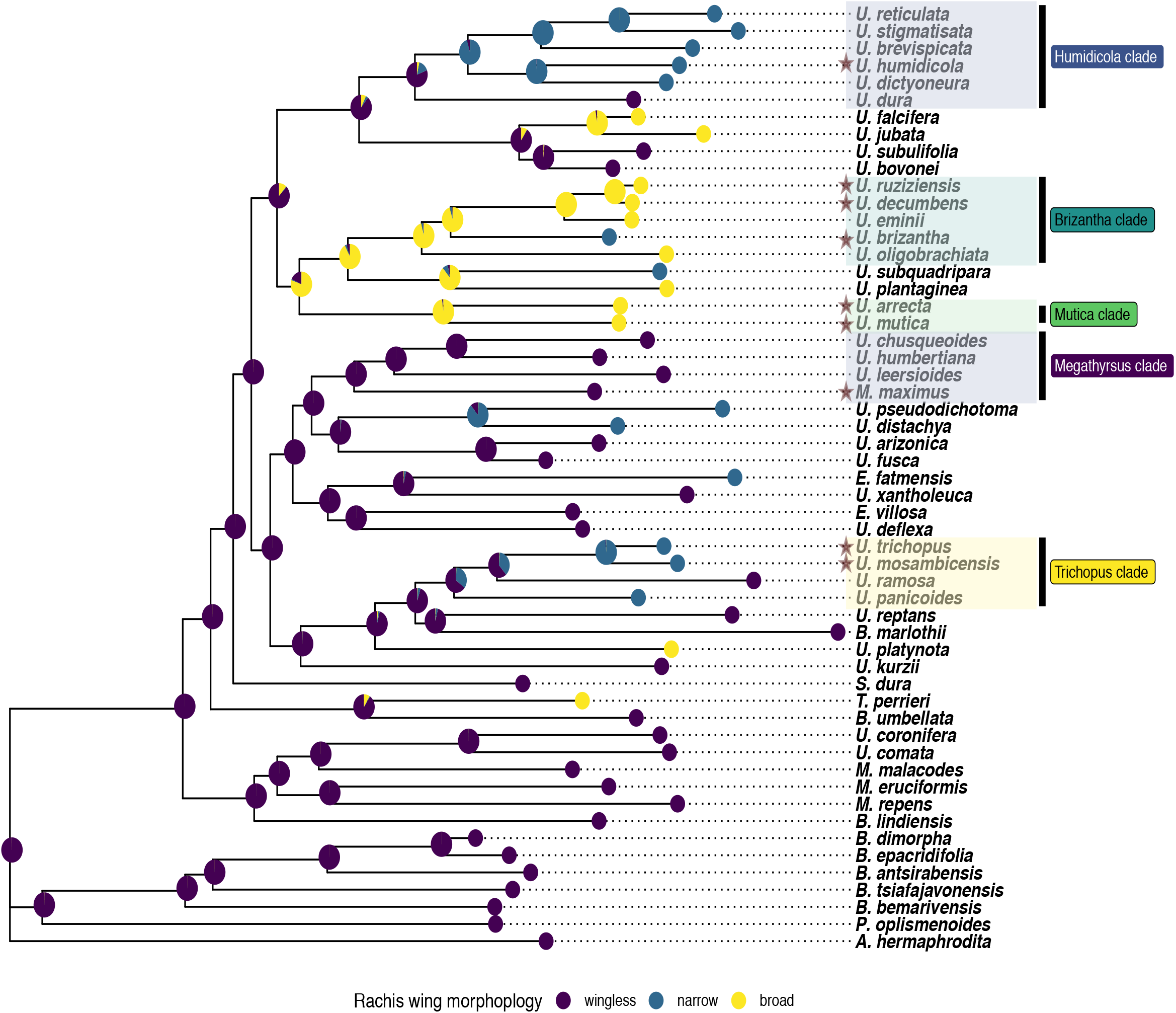
Evolution of the rachis morphology. Maximum likelihood tree estimated using IQTREE2 v2.1.2 inferred from 327 nuclear genes. Ancestral state estimations for an rachis morphology were performed using corHMM in R and posterior probabilities for state estimations mapped to ancestral nodes.

**Table 3:** Model selection for growth habit (annual vs perennial) and rachis wing morphology with log likelihood, Akaike information criterion (AIC), and AIC delta scores.

| Trait | model | log likelihood | AIC | delta.AIC |
| --- | --- | --- | --- | --- |
| Growth Habit | Equal Rates | -43.0951 | 88.19 | 0 |
|  | All Rates Different | -43.0951 | 90.19 | 2 |
|  | Annual to Perennial | -46.12271 | 94.245 | 6.055209 |
|  | Perennial to Annual | -49.5135 | 101.03 | 12.8368 |
| Rachis morphology | Equal Rates | -51.12791 | 104.26 | 0 |
|  | All Rates Different | -47.87176 | 119.74 | 15.4877 |
|  | Symmetrical Rates | -48.81644 | 109.63 | 5.377079 |

As the *Brizantha* and *Mutica* clades are the most closely related forage clades, posterior probabilities indicate that the shared ancestor of these important forage clades may have already evolved a broad rachis wing. For the *Humidicola* clade, results indicate that the clade likely evolved from a wingless ancestor, but that a narrowly winged form likely emerged at an early point in the clade’s divergence, as the vast majority of *Humidicola* clade species possess winged rachises.

## Discussion

### Crop Wild Relatives for *Urochloa* forage breeding

The phylogenetic analyses conducted here provide a platform for interpreting the evolution of *Urochloa s.l.*. A broad definition of *Urochloa s.l.* (which includes all subsumed genera) is support in both trees, confirming previous results based on chloroplast markers (Salariato et al., 2010, 2012; Hackel et al., 2018; Delfini et al., 2023). However, inferring trees using hundreds of nuclear loci allowed us to resolve polytomies previously seen in *Urochloa s.l.* phylogenies inferred using a handful of chloroplast markers. Further, both ASTRAL-III and ML trees recovered topologies more congruent with, albeit informal, analyses of infrageneric groupings in *Urochloa* (Renvoize et al., 1996). For example, Salariato et al. (2010;2012) and Delfini et al. (2023) consistently fail to recover the *Humidicola* and *Mutica* clade using chloroplast markers. Congruence between our phylogenetic analyses indicates that they represent a more realistic model of *Urochloa* diversification.

For all forage species, we were able to identify wild grasses that are plausible CWRs based on monophyly in both phylogenetic analyses. CWRs are important taxa for agricultural purposes as they provide the comparative context for more in depth analysis for phenotypic developmental and molecular studies in crops (Harlan and DeWet, 1971; Pironon et al., 2020; Viruel et al., 2021). Additionally, crosses between crop species and their wild relatives can produce cultivars with improved desirable traits, e.g. fungal and nematode disease resistance in peanuts (Bertioli et al., 2021), and increase heat and drought tolerance in wheat (Molero et al., 2023). Interspecific crossing has become routine in *Urochloa* cultivar production as the most popular commercially available lines, Mulato and Mulato II, were developed from *U. brizantha* x *U. decumbens* x *U. ruziziensis* hybrids (Argel et al., 2007; Pizarro et al., 2013).

*Urochloa brizantha*, *U. decumbens*, and *U. ruziziensis* share a recent common ancestor with wild relatives *U. eminii* and *U. oligobrachiata*, confirming the observations of Renvoize et al. (1996) based on spikelet and inflorescence morphology. Further, alpha taxonomists have long argued that intermediate specimens are common between *U. ruziziensis* and *U. decumbens*, and *U. decumbens* and *U. brizantha* respectively (Renvoize and Maass, 1993; Renvoize et al., 1996; Sosef, 2016), demonstrating that detailed morphological studies still provide insight into evolutionary dynamics even in highly intricate species complexes. Sosef 2016 argued that *U. decumbens* and *U. ruziziensis* be lumped together along with *U. eminii,* and our analysis confirms that the three taxa are the most closely related within the *Brizantha* complex. It is likely that specimens identified as *U. eminii* are not used in cultivar breeding programs. *Urochloa oligobrachiata*, a perennial diploid species with large leaves and seeds within the *Brizantha* clade, is also absent in breeding programs and is not represented in global germplasm banks (to our knowledge). It is plausible that both species could be successfully integrated into breeding programs, and the genetic and phenotypic contributions this could make towards cultivar production should be explored.

Cultivar development for *U. humidicola* forages lags behind the *Brizantha* clade at a commercial scale (Boldrini et al., 2011). While apomictic reproduction in the complex has been overcome by the chance discovery of sexual reproductive accessions (Jungmann et al., 2009), the hexaploid nature of the complex and reduced genetic diversity in seed bank collections creates a bottleneck for breeders (Miles, 1996; Vigna et al., 2016, Higgins et al., 2022). Introducing CWR to breeding systems remains a viable option for overcoming these limitations. Contrasting our molecular results with the morphological analysis of Renvoize et al. (1996) provides more useful insight into the Humidicola clade diversification. Renvoize et al. (1996) also identified *U. stigmatisata*, *U. reticulata*, and *U. brevispicata* as close relatives of *U. humidicola* and *U. dictyoneura*. However, the authors grouped *U. dura* with *Brizantha* clade species, whereas our analysis shows it belongs within the *Humidicola* clade. Further, Renvoize et al. (1996) placed *U. platynota* as a close relative to the *Humidicola* clade, but our analysis shows it is a distantly related species. Finally, the authors found that *U. falcifera*, *U. jubata*, *U. subulifolia*, and *U. bovonei* shared were closely related to *Humidicola* clade species. Our analysis shows these taxa form a well-supported clade sister to the *Humidicola* clade.

Broadly, a suite of CWR and sister taxa to the *U. humidicola/U. dictyoneura* complex have been inferred in the literature and strongly supported by our phylogenies. Introducing CWR and interspecific breeding to *U. humidicola*/*U. dictyoneura* could bring cultivar production up to speed with *Brizantha* clade cultivars. Additionally, cytological evidence and fluorescent in-situ hybridization have provided evidence for the inferred allopolyploid origins of *U. humidicola* and, crucially, potential subgenome identification in the species (Vigna et al., 2016; Tomaszewska et al., 2023). CWRs provide a sensible starting point for investigating the putative donors of *U. humidicola* subgenomes, as has been demonstrated across numerous allopolyploid crop species (He et al., 2017; Edgar et al., 2019; Yim et al., 2022).

*Trichopus* and *Mutica* clade species are less commercially important in a global context, though their importance as livestock feed at small scales has been noted (Fischer and Kerridge, 1996; Pereira Filho et al., 2013). *Urochloa trichopus* and *U. mosambicensis* have been shown to be a nutrient dense food source for goats in low-precipitation regions of Brazil such as the Caatinga (Pessoa et al., 2022). While the *Trichopus* clade is distantly related to other forage clades, the *Mutica* clade shares a recent common ancestor with the *Brizantha* clade. The placement of *Megathyrsus* within *Urochloa* is strongly supported, and its placement with species with more broadly spaced and pedicelled spikelets i.e. *U. chusqueoides* and *U. humbertiana* (Renvoize et al., 1996) provides a sensible framework for comparative analysis of inflorescence diversity in *Urochloa s.l*.

### Agricultural trait evolution

Modelling character evolution is challenging in groups where data availability is limited for both taxa and appropriate traits. Herbarium accessions play a pivotal role in bolstering taxon representation in phylogenies, especially for groups with geographical ranges spanning multiple countries and continents such as *Urochloa* (Besnard et al., 2014; Baker et al., 2021, Larson et al., 2023). Comprehensive phylogenies are commonly leveraged for evolutionary and ecological trait comparisons in numerous biological disciplines (Revell and Harmon, 2022), but the application of these methods in assessing agricultural potential across plant species (and clades) is underexplored. Forage grasses present a unique opportunity to apply phylogenetic comparative methods for agricultural purposes as traits of interest for breeders, taxonomists, and ecologists share considerable overlap (leaf size, plant height, growth habit, etc.) and are likely present in floras and databases (Clayton and Renvoize, 1982; Clayton et al., 2016). For select *Urochloa* species, the domestication process is well within the initial phases (Dusi et al., 2010; Jank et al., 2014).

High and statistically significant phylogenetic signals for all three continuous traits assessed in this study imply that, assuming a Brownian Motion model of evolution, leaf area, seed size, and culm height are shared across *Urochloa* species due to their shared history (Revell et al., 2008). Congruence between Blomberg’s K and Pagel’s **λ** (values approach 1 for all three traits using both approaches) provide strong support for phylogenetic signal in leaf and seed size in *Urochloa*. Ancestral trait estimates for continuous traits generally show size increases along branches from ancestral clade nodes for important forage species and their close relatives. Our discrete character state estimations show a similar trend. A winged rachis morphology emerged independently in all forage clades except for the *Megathyrsus* clade where all species (including *M. maximus*) have wingless rachises and lax inflorescences. A winged rachis has been noted to impose greater rigidity on spikelet ordering (Renvoize et al., 1996), and is partly associated with a ‘homogenised’ inflorescences morphology as outlined by Salariato et al. (2010) and Reinheirmer and Vegetti (2008). Further investigation of how inflorescence structure influences seed retention (non-shattering phenotype) in *Urochloa* essential, as non-shattering is among the first selected traits in plant domestication for grain production (Konishi et al., 2006; Yu et al., 2020). Estimating ancestral growth habit shows more state uncertainty, particularly at deeper nodes in the phylogeny. However, there remains evidence that perennial growth habits evolve multiple times across *Urochloa* in forage clades. Improving certainty for node state estimates can be achieved by more dense sampling of *Urochloa* species in future.

Palatable perennial grasses are common across the African continent (Ezenwa et al., 2006; Bond 2010), though the independent emergence of species with forage potential across *Urochloa* is notable. The goal of forage grass breeding is to develop cultivars with unique phenotypes to suit specific geographical and climatic regions, while not sacrificing nutritional content and biomass production (do Valle, 2013; Nguku, 2015). Achieving this goal sustainably will require selecting material from genetically diverse accessions and taxa (Ferreira et al., 2021). Based on our results, the independent evolution of forage syndromes across *Urochloa* implies a high amount of taxonomic and genetic diversity that forage breeders can draw from for future cultivar development. Therefore, *Urochloa s.l.* conservation across its natural distribution, and particularly in Africa, should be a high priority for commercial breeders.

### Future considerations

The forage potential for *Urochloa* at the genus level clearly outmatches cultivar production across the tropics. Introducing CWRs into future breeding programs is a steppingstone towards improving commercially available grasses. Interspecific *Urochloa* crosses are only successful if ploidy levels between parental species match and at least one parent is sexually reproductive. Addressing these knowledge gaps will require high-quality, chromosome scale genome assemblies for important forage species and CWR (Risso-Pascotto et al., 2005; Simeão et al., 2021). Within the forage clades identified in this study, only chromosome-scale genome assemblies exist for *U. ruziziensis* (Pessoa-Filho et al., 2019, available online but analysis unpublished; Worthington et al., 2021). Additional high-quality genome assemblies will be valuable and can be used to determine ancestral genome origins in polyploids, chromosome rearrangements in the various species with distinct chromosome numbers and provide reference genomes for alignment of polymorphic markers from reduced-representation sequencing.

Genome assemblies could reveal the genetic pathways associated with *Urochloa* invasiveness into non-agricultural land, an unfortunate trend seen in African grasses globally (Visser et al., 2016). For example, *Mutica* clade species have been introduced from African countries to various parts of the world with the putative aim of improving pastures for livestock rearing (Williams and Baruch, 2000). While these species clearly have good forage characteristics, demonstrated in this study and elsewhere (Fischer and Kerridge, 1996; Veldkamp 1996), little scientifically informed breeding has been attempted in *U. mutica* and *U. arrecta*, and the two species are commonly classified as invasive weeds outside the African continent (Boyden et al., 2019). Even in the commercially important *Brizantha* clade, *U. decumbens* is an aggressive invasive in the Cerrado, a dry savannah region in Brazil (Pivello et al., 1999). This is likely a consequence of the species’ early introduction to South America as a forage grass prior to the establishment of genetic breeding programs (Pivello et al., 1999; Barbosa et al., 2008). There exists a substantive link between forage potential and aggressive invasiveness in African grasses, and genomic resources could help mitigated this undesirable attribute (Daehler and Carino, 1998; Williams and Baruch, 2000; Cook and Dias, 2006; Barbosa et al., 2008; Barbosa, 2016).

Beyond this study, there is still a need for more in-depth knowledge of the basic biology and diversity in *Urochloa s.l.*, and greater emphasis must be placed on conserving and collecting wild *Urochloa* grasses, particularly in African countries. While commercial cultivars are predominantly utilised at large scale in South America (Jank et al., 2014; Maass et al., 2015), African nations have begun reintroducing cultivars in beef, dairy, and push-pull pest control systems with notable successes (Mutimura and Everson, 2012; Khan et al., 2014; Clémence-Aggy et al., 2021). As the centre of *Urochloa s.l.* diversity, natural populations of African species likely contain the genes and traits needed to tailor new cultivars for the specific and varying needs of farmers, livestock, and ecosystems across African nations. Conservation of African grasses is a global sustainability imperative as African grasslands form the basis of ancient habitats (Bond, 2016; Solofondranohatra et al., 2020; Buisson et al., 2022), perform natural carbon sequestration (Vågen et al., 2005; Dobson et al., 2022), and support the livelihoods of millions of people and animals (Bengtsson et al., 2019). African grassland conservation safeguards the biodiversity needed to address issues of economic development and food security, and *Urochloa* is a genus of primary consideration in this regard.

## Conclusion

We have reconstructed a nuclear phylogeny for the grass genus *Urochloa s.l.* that is both comprehensively sampled and data rich, focusing on forages and their relatives. Our phylogenomic analysis allowed us to infer the placement of key agricultural species within the genus and identify their closest wild relatives. Additionally, we were able to estimate the ancestral state of numerous agriculturally important traits and demonstrate their convergent emergence in agriculturally important lineages. *Urochloa s.l.* is a highly morphologically diverse genus replete with polyploidization events and a natural distribution spanning the near entirety of the southern hemisphere. These attributes make *Urochloa* a prime example of how African grasses should serve as model systems for studying complicated evolutionary events, how a strong taxonomic and phylogenetic foundation can aid these studies, and how this knowledge can facilitate more sustainable agricultural practices in countries where it is most required.

## Supplementary data

Supplementary data are available online at https://academic.oup.com/aob and consists of the following: Table 1: Specimen data with trait data from GrassBase (Clayton et al., 2016), Table 2: Forage clades, CWR, and forage traits, Figure 1: Ancestral trait estimation for leaf area, Figure 2: Ancestral trait estimation for culm height, Figure 3: Ancestral trait estimation for fertile lemma length

## Acknowledgements

The authors acknowledge contributions from Jan Hackel, Philipps-Univesität Marburg, for providing sequence read data, and the James Hutton Institute and NIAB for proving computational resources through the ‘UK’s Crop Diversity Bioinformatics HPC” (BBSRC grant BB/S019669/1) which contributed to the results obtained in this paper.

