## Supplementary Figure 3 for "Phylogenomic analysis reveals the evolutionary origins of five independent clades of forage grasses within the African genus *Urochloa*"

Masters et al. 2023

Supplementary figure 3

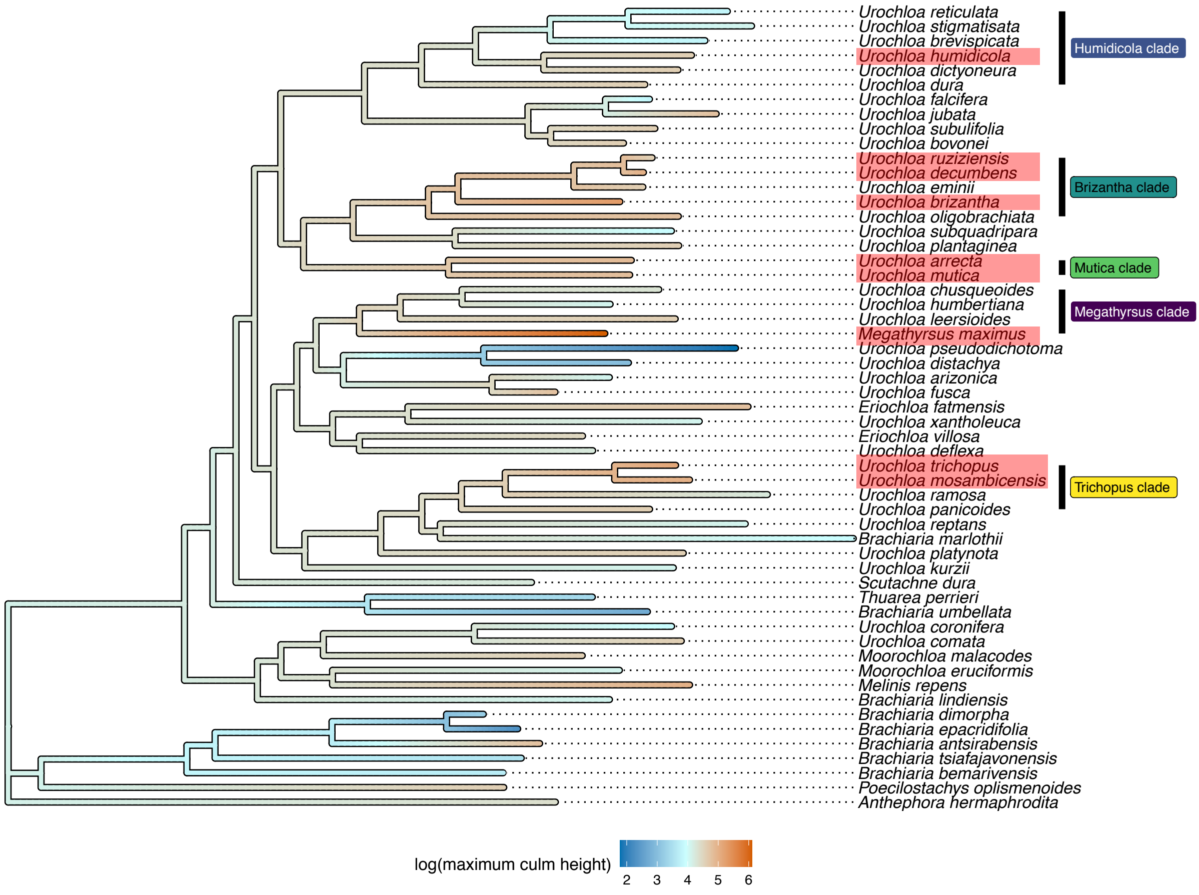


Evolution of culm height. Maximum likelihood tree estimated using IQTREE2 v2.1.2 inferred from 327 nuclear genes. Log transformed maximum culm height (cm) in *Urochloa s.l.*. Ancestral trait estimation along branch lengths.
