## Supplementary Figure 2 for "Phylogenomic analysis reveals the evolutionary origins of five independent clades of forage grasses within the African genus *Urochloa*"

Masters et al. 2023

Supplementary figure 2

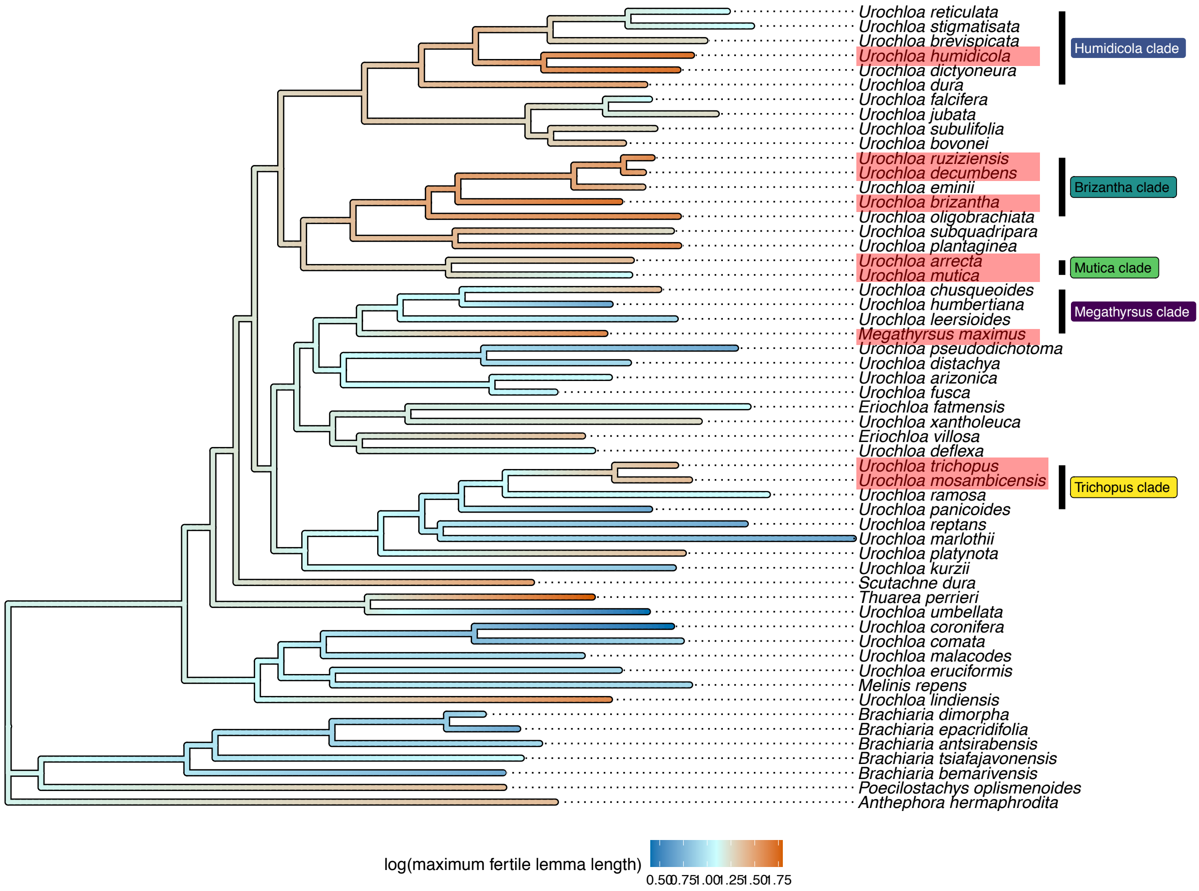


Evolution of lemma length. Maximum likelihood tree estimated using IQTREE2 v2.1.2 inferred from 327 nuclear genes. Log transformed maximum fertile lemma length (mm) in *Urochloa*. Ancestral trait estimation along branch lengths
