## Supplementary Figure 1 for "Phylogenomic analysis reveals the evolutionary origins of five independent clades of forage grasses within the African genus *Urochloa*"

Masters et al. 2023

### Supplementary figure 1

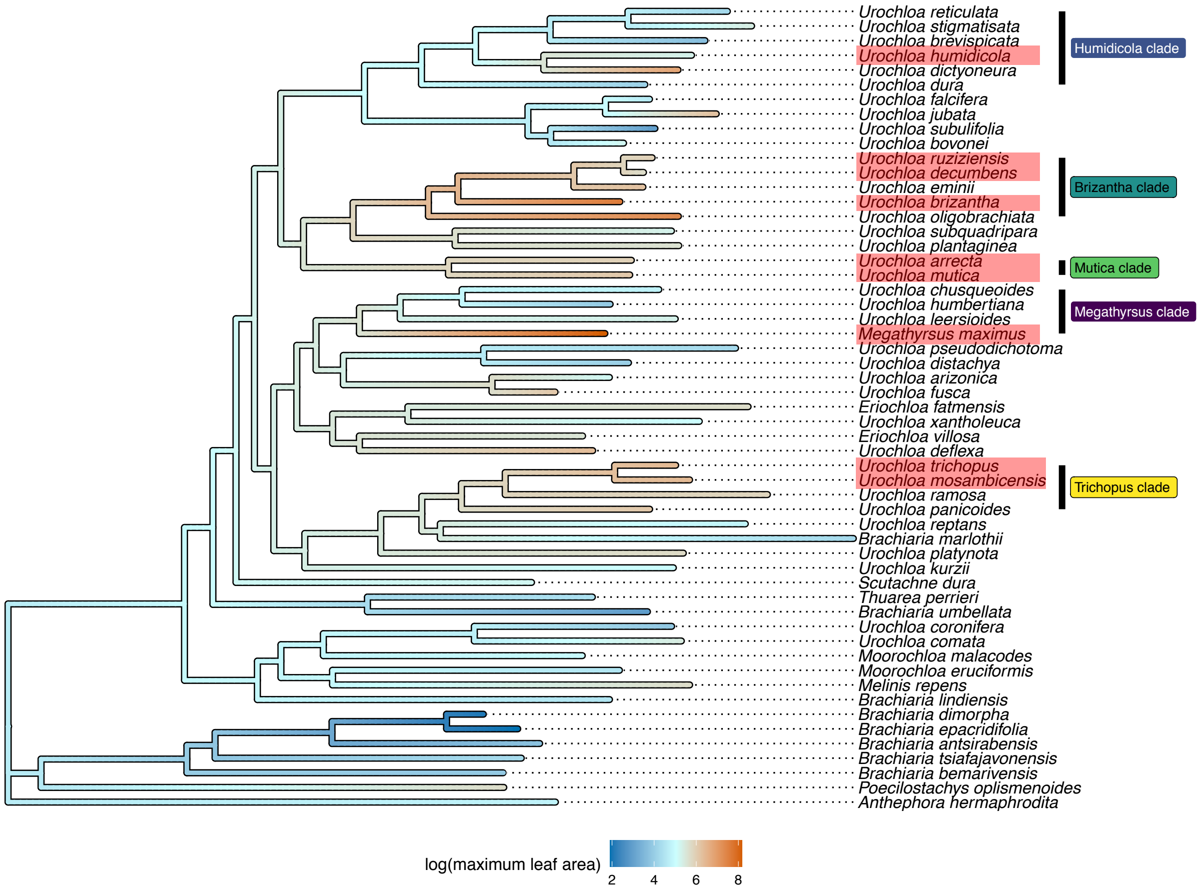


Evolution of leaf area. Maximum likelihood tree estimated using IQTREE2 v2.1.2 inferred from 327 nuclear genes. Log transformed maximum leaf area ($cm^2$) in *Urochloa.* Ancestral state estimations along branch lengths.
